# Monitoring the dynamics of exosomes through the tracking of Rab7 partners

**DOI:** 10.1101/2025.10.07.680694

**Authors:** Yonis Bare, François Delalande, Aurélie Hirschler, Chantal Cazevieille, Vincent Lucansky, Christine Carapito, Raphael Gaudin, Maika S. Deffieu

## Abstract

Extracellular vesicles (EVs) play a crucial role in cell-cell signaling, and their dysregulation is linked to various pathologies. EVs can originate from the plasma membrane as microvesicles, or form within endosomal compartments that fuse with the plasma membrane, releasing their intraluminal vesicles in the extracellular space as exosomes. However, distinguishing between these two types of EVs is challenging due to their common protein markers, necessitating cumbersome isolation methods. To overcome this limitation, we developed an innovative tool using Rab7-mediated proximity labelling and mass spectrometry. This technological advancement enables the covalent labeling of Rab7 partners allowing the tracking of intracellular and extracellular exosomal cargoes.

## Introduction

Extracellular vesicles (EVs) are small membranous vesicles secreted by various cell types. They mediate intercellular communication^1^, can have a function in immune responses^2^, carry viruses to favor their spreading^3^, and participate in tumor progression by modifying the tumor micro-environment^4^. EVs are an heterogenous population with different subtypes of vesicles that can carry various cargoes such as lipids, nucleic acids, and proteins^5,6^. While current isolation methods can distinguish EV subtypes based on size, density and other biophysical characteristics, they do not allow us to distinguish between their subcellular sites of biogenesis^7^.

However, two main types of vesicles were described: microvesicles, which originate from the plasma membrane, and exosomes, which originate from multivesicular bodies (MVBs). Microvesicles have a size between 50-1000 nm and are formed by the assembly of specific cargoes at the plasma membrane, followed by their outward membrane budding to shape a vesicle, which will be released by fission from the plasma membrane^8,9^. Exosomes have a size between 50-150nm and are released by the fusion of MVBs containing intraluminal vesicles (ILVs) with the cell surface. Cargoes are packaged into ILVs which are formed by an inward budding of the limiting endosomal membrane and then released into the endosomal lumen^10^. The biogenesis of ILVs can influence the specificity of cargo loading, potentially involving different molecular mechanisms that are still not fully understood. Previous work has demonstrated that ILV biogenesis involves the endosomal sorting complexes required for transport (ESCRT) machinery. This machinery participates in various stages of ILV formation: ESCRT-0 is involved in cargo selection, ESCRT-I and ESCRT-II in membrane deformation, and ESCRT-III in membrane scission^11,12^.

ILV biogenesis could be formed through other ESCRT-independent pathways involving the type II sphingomyelinase in oligodendrocytic cells^13^. Tetraspanins, such as CD63, also participate in cargo sorting in melanocytic cells^14^. Additionally, Flotillin-1 and Flotillin-2 (Flot1 and Flot2 respectively), which mainly function as scaffold proteins, have been suggested to be important for EGFR entry inside exosomes^15^. Recently, it was demonstrated that cytosolic proteins containing the KFERQ motif were loaded by Lamp2A into a subtype of small EVs^16^. The identification of the different subpopulations of EVs remains unclear as their formation depends on the cell type, or pathological state of the cell.

To distinguish between EV subtypes, dedicated techniques were developed to determine their respective protein content. These techniques involved purification methods such as differential ultracentrifugation, filtration, or density gradients to separate the different populations of EVs. Proteomic studies were then conducted to identify specific EV markers, as these markers can vary depending on the cell type^5,17^. Despite these improvements, it remains challenging to differentiate between the various EV subpopulations due to their shared protein markers, including the tetraspanins CD9, CD81, CD63, and the scaffold proteins Flot1 and Flot2.

To gain a deeper understanding of EV biology, researchers have been investigating the tracking of these vesicles from their biogenesis, transport, and subsequent secretion out of the cell. Rab-GTPases, which are essential regulators of membrane trafficking, play a key role in the formation of these vesicles. Rab27A and Rab27B are involved in exosome secretion, mediating the trafficking and docking of MVBs to the plasma membrane^18^. Rab7A, which traffics from late endosomes to lysosomes, was found to be involved in exosome secretion upstream of the fusion events between MVBs and the plasma membrane^19,20^. A recent study implies ER-membrane contact sites in the formation of CD63-positive exosomes. Their secretion depends on their transport through sequential switches between Rab7A, Arl8B and Rab27B-positive vesicles before reaching the plasma membrane^19^. To track the path of these vesicles, genetic engineering of the EV marker CD63 fused to pHluorin was used to monitor MVB fusion with the plasma membrane and the release of vesicles in the extracellular environment^19^. Another recent study indicated that Rab31 is involved in the secretion of small EVs containing the KFERQ motif^16^.

In our study, we employed a proximity labeling strategy to monitor EVs generated directly from MVBs, i.e. exosomes. We utilized the engineered ascorbate peroxidase 2 enzyme (Apex2) as a proximity labeling assay method to temporally biotinylate all protein interactors surrounding the protein of interest within a radius of 20nm ^21,22^. The reaction is activated by hydrogen peroxide, and biotin-tyramide is used as a substrate. The biotin-tyramide radicals formed after catalysis are short-lived, leading to the specific crosslinking of biotin at exposed tyrosine residues of nearby proteins. To specifically monitor EVs originating from endosomes, we used the MVB-localized protein Rab7A fused to the Apex2 enzyme (Apex2-Rab7A). We combined quantitative proteomics with the proximity labeling assay to identify EV markers labeled by Apex2-Rab7A. Among the relevant candidates, we identified Flot1 and Flot2 as exosome cargoes. Interestingly, we demonstrated that Rab7A can interact with Flot1. Moreover, Rab7A expression could modulate the loading of Flot1 in small EVs and large EVs. A more precise characterization of these isolated EVs revealed the presence of biotinylated cargoes and endogenous Rab7A, supporting the endosomal origin of the biotinylated cargoes. Altogether, our tool suggests a new function of Rab7A in EV cargo loading and enables the specific labeling of exosomes intracellularly.

## Results

### Flag-Apex2-Rab constructs are active and properly localized in cells

Here, we used a proximity labeling assay to label Rab7A potential partners in a spatio-temporal manner. We generated Flag-Apex2-Rab7A and two other Rabs that are used as control of specificity (Flag-Apex2-Rab5C and Flag-Apex2-Rab6B). These constructs were placed under the control of the promoter of the mouse gene encoding the ribosomal protein L30 for weak mRNA and protein expression, which favors the proper localization of Rab proteins. Rab7A, Rab5C, and Rab6B proteins were fused at their N-terminus with a Flag peptide for protein detection followed by the Apex2 for partners labeling (Fig. 1A). Upon transfection of SUM159 cells, immunofluorescence analysis indicated a correct localization of Flag-Apex2-Rab7A, Flag-Apex2-Rab5C, and Flag-Apex2-Rab6B proteins. Flag-Apex2-Rab7A exhibited a cytosolic pattern as well as a vesicular profile which localized with Lamp1 positive vesicles, an endo-lysosomal marker (Fig. 1B, left panel, white arrows). Flag-Apex2-Rab5C was also expressed in the cytosol and in vesicles positive for the Rab5 effector, EEA1, localized in early endosomes (Fig. 1B, middle panel, white arrows). Flag-Apex2-Rab6B was present at steady-state at the Golgi apparatus and colocalized with the trans Golgi marker TGN46 (Fig. 1B, right panel, white arrows). Together, these data indicate that the addition of the Flag tag and the Apex2 protein does not disturb the localization of these small Rab-GTPases. To determine if the Apex2 is functional, we monitored the distribution of the biotinylated proteins using an anti-biotin antibody. Cells were placed in the presence of H_2_O_2_ to activate the Apex2, and biotin-tyramide was added for biotinylation of proximity partners (see Material & Methods for details). We observed again a cytosolic and vesicular pattern for the localization of Flag-Apex2-Rab5C and Flag-Apex2-Rab7A, while the biotin staining was exclusively found in vesicles (Fig. 1C, white arrows). In the case of Flag-Apex2-Rab6B, the biotinylated partners were secluded to the Golgi apparatus, which is the usual subcellular localization of Rab6B at steady state (Fig. 1C). Western blot analysis showed that biotinylated partners, enriched with streptavidin resin, were detected solely in the presence of H_2_O_2_ (Fig. 1D), suggesting that the biotin labeling is specific to Apex2 activation. To discriminate between endosomal and plasma membrane-derived EVs, Flag-Apex2-Rab7A must be strictly localized at the endosomes and not at the plasma membrane. To ensure its presence at the endosomes, we used transmission electron microscopy in presence of 3,3′diaminobenzidine tetrahydrochloride (DAB). The DAB reacts with the Apex2 enzyme, and the product formed precipitates locally to become dense to electrons following osmium staining. As expected, we observed the presence of intracellular vesicles that are dense to electrons in presence of DAB compared to the control without DAB. In contrast, no DAB staining was visible at the plasma membrane indicating the absence of Flag-Apex2-Rab7A at that location (Suppl. Fig. S1, white arrows). Together, we showed that the engineered Flag-Apex2-Rab constructs are functional and distribute at expected locations.

**Figure 1:**
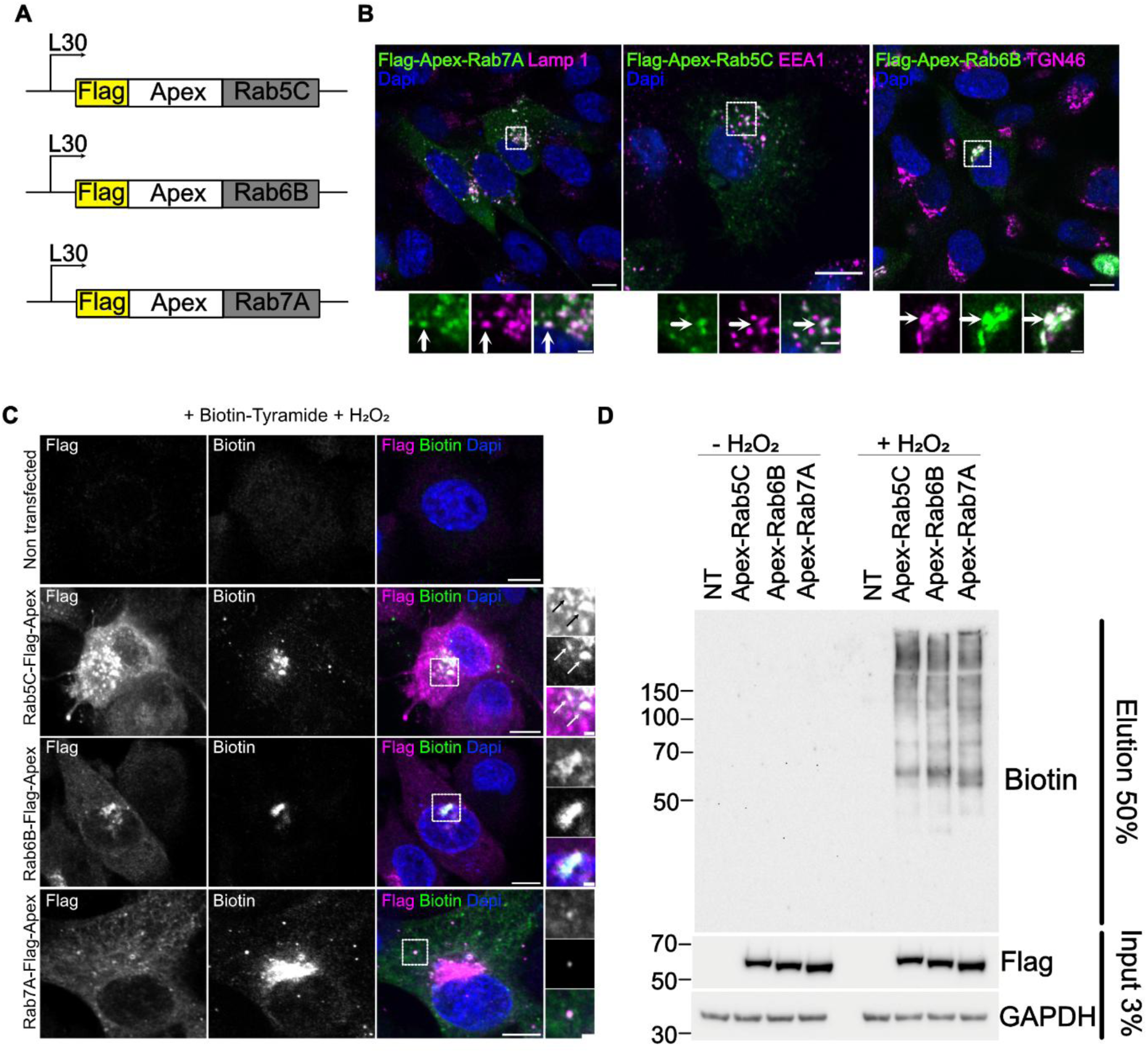
Flag-Apex-Rab proteins are properly localized and active. (A) Schematic on the Flag-Apex2-Rab constructs. L30 is used as a weak promoter, Flag and Apex2 inserted at the 5’ of the Rab7A, Rab5C and Rab6B cDNA sequence. The size of the boxes is not to scale of the number of nucleotides. (B) Confocal images indicating Flag-Apex2-Rabs localization. Flag-Apex2-Rab7A, Flag-Apex2-Rab5C, Flag-Apex2-Rab6B were expressed in SUM159 cells. After 24h post-transfection cells were fixed and stained. Flag-Apex2-Rab constructs were visualized with Flag antibody (green). Antibodies against Lamp1 (magenta) labeled the endo-lysosomal compartments, antibodies against EEA1 (magenta) were used for early endosomes, and antibodies against TGN46 (magenta) labeled the trans-Golgi compartment. A representative image from a single z-stack was represented, zoomed images were shown in lower panels. The co-distribution of Flag-Apex2-Rab proteins with Lamp1, EEA1 or TGN46 were indicated with white arrows. Scale bars = 10 µm; scale bars for zoomed images = 2 µm. (C) Confocal images visualizing Apex2 activity. SUM159 cells were non-transfected (NT) or transfected with Flag-Apex2-Rab7A, Flag-Apex2-Rab5C, Flag-Apex2-Rab6B. After 24h post-transfection, Apex2 was activated with 500µM hydrogen peroxide. Protein biotinylation was visualized using an antibody against biotin (green), Flag-Apex2-Rabs were observed with an antibody against the Flag tag (magenta), Dapi was used to label the nuclei (blue). Co-distribution of Flag-Apex2-Rab proteins with biotinylated proteins were visualized in zoomed panels. Scale bars = 10 µm, scale bars for zoomed panels = 2µm. (D) Western blot analysis demonstrating the presence of biotinylated proteins. Cells were transfected with Flag-Apex2-Rab7A, Flag-Apex2-Rab5C, Flag-Apex2-Rab6B and were placed in presence or in absence of 2 mM H_2_O_2_ at 24 h post-transfection. Streptavidin affinity chromatography was performed to enriched biotinylated protein (elution). Biotinylated proteins were visualized with a biotin antibody. Input samples indicated the expression of the Flag constructs observed with a Flag antibody, and anti-GAPDH antibody was used as a loading control.

As biotinylation cannot be observed in live cells, we next applied the proximity labeling assay technique for live imaging studies using the cell-permeant tetramethylrhodamine-tyramide (TMR-tyr) as a substrate of the Apex2 enzyme. TMR-tyr was successfully associated to vesicles positive for Flag-Apex2-Rab7A following H_2_O_2_ activation in fixed cells (Fig. 2A). To confirm the specificity of the TMR labelling, we built a novel ALFA-tagged construct (referred to as ALFA-Apex2-Rab7A), a 13 amino acid tag, which can be recognized by a genetically-encoded nanobody fused to a fluorescent protein^23^. SUM159 cells stably expressing miRFP680-nanobody and transfected with ALFA-Apex2-Rab7A revealed that this construct localizes at the endosome as confirmed by the co-distribution with EGFP-Rab7A. The TMR-tyr labeling was associated to Rab7A positive vesicles (Fig. 2B, upper panel, white arrows) and CD63-EGFP, a traditional MVB marker (Fig. 2B, lower panel, white arrows). This result highlights that our method can be used for live imaging of Rab7A and its close partners.

**Figure 2:**
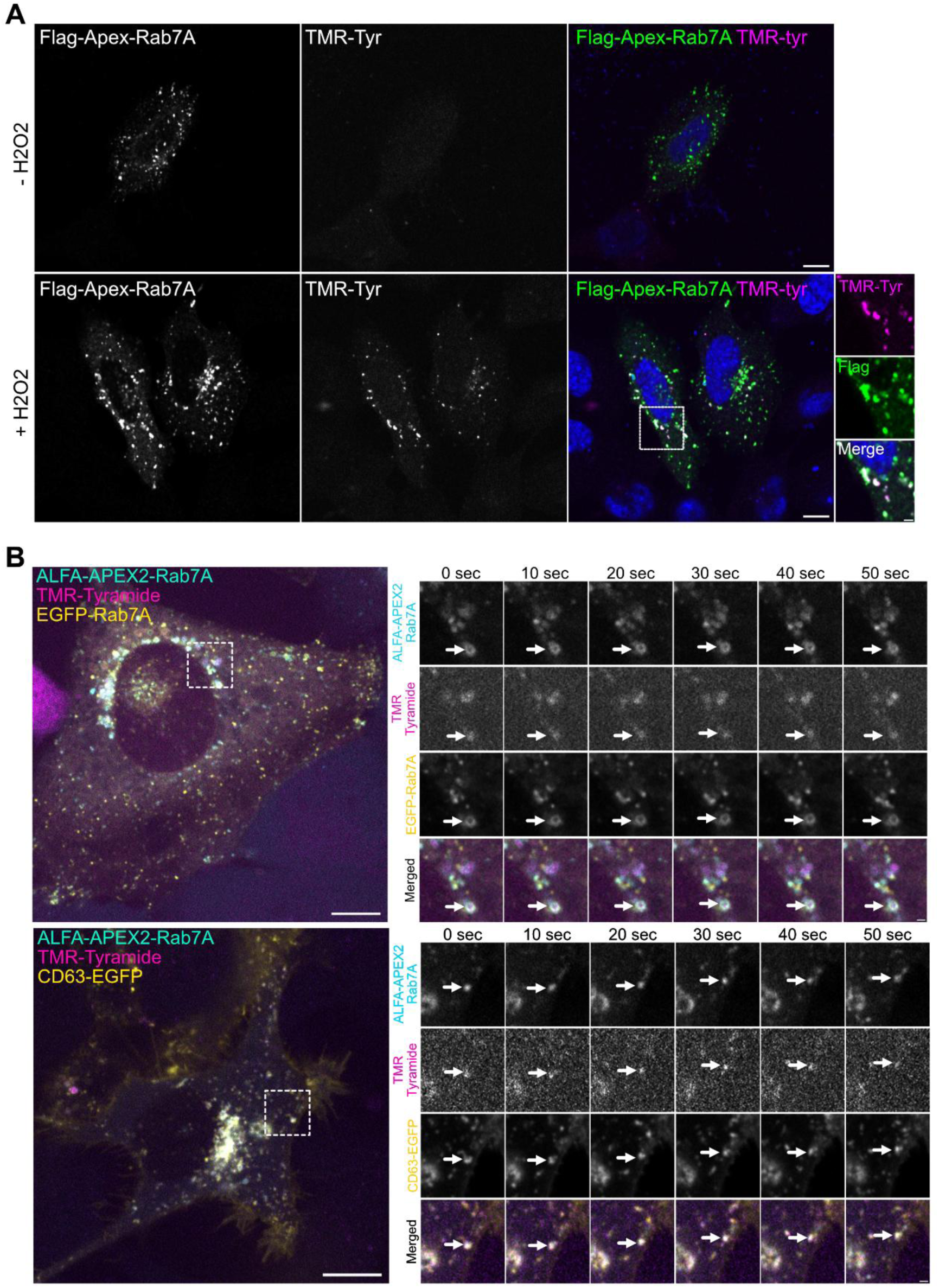
Fluorescent proximity labelling of partners involved with Rab7A. (A) Cells were transfected with Flag-Apex2-Rab7A for 24 h and incubated with or without 500 µm of hydrogen peroxide. TMR-Tyramide (TMR-Tyr) was visualized in magenta and Flag-Apex2-Rab7A was visualized with anti-Flag antibody (green) Scale bar = 10 µm, scale bars from zoomed regions (dotted box) = 2 µm. (B) Representative still images from videos taken of SUM159 cells stably expressing Nanobody fused to miRF680 fluorescent protein (cyan) and transfected with plasmids encoding ALFA-tagged Apex2-Rab7A and EGFP-Rab7A, or CD63-EGFP (yellow). Proximity partners of Apex2-Rab7A were labelled using a tyramide group coupled to tetramethyl rhodamine (TMR Tyramide, magenta). Left panel represents still image from video at t = 0 s, scale bar = 10 µm. Right panel represents still images from zoomed inset (dotted box), scale bar = 1 µm.

### Quantitative mass spectrometry analysis reveals EV markers as Rab7A molecular partners

To identify the proteins associated with Rab7A, we combined the PLA approach with mass spectrometry analysis. We performed the PLA in SUM159 cells in the presence of biotin-tyramide with Flag-Apex2-Rab7A as the bait of interest. Non-transfected and Flag-Apex2-Rab5C controls were used to select specific biotinylated partners. Cells were incubated with biotin alone (that cannot be covalently linked) or with biotin-tyramide (the substrate of Apex2). Cells were lysed in the presence of antioxidants and biotinylated proteins were purified with streptavidin affinity chromatography resins (Fig. 3A). Western blot analysis confirmed the presence of biotinylated proteins in the Flag-Apex2-Rab5C and Flag-Apex2-Rab7A conditions (Fig. 3B). Although a prominent non-specific band was seen in the non-transfected (NT) and the transfected conditions, a smear of biotinylated proteins indicated the presence of potential hits. Eluted samples were submitted to nanoLC-MS/MS and differential proteomic analyses were performed. We identified more than 700 proteins for each condition (Fig. 3C). To select the most promising candidates, two criteria were applied: proteins above a fold change of 1.5 and above a p-value (significance -Log_10_) of 2 were considered. Non-transfected cells (NT) with biotin-tyramide were used as a control for non-specific biotinylation (Fig. 3D). To select specifically Rab7A partners, the comparison was also performed with cells expressing Flag-Apex2-Rab5C in the presence of biotin-tyramide (Fig. 3E). To remove proteins that were not biotinylated and might bind non-specifically to the agarose beads, a background control Flag-Apex2-Rab7A with biotin alone was also performed (Fig. 3F). Summarizing these data, the selected proteins were subjected to gene ontology analysis via the g:Profiler tool^24^ and investigations on the cellular components were represented (Suppl. Fig. S2A). The enriched Rab7A partners were significantly associated to pathways such as focal adhesion, cortical cytoskeleton, and extracellular vesicles (-log_10_ p-value above 2; Suppl. Fig. S2B). The most enriched hits were Rab7A, which could be auto biotinylated by Apex2, and Flot1 and Flot2 (Table 1 and Suppl. Fig. 2A-B), which are commonly found in EVs^17^. Annexin A7 (ANXA7) was also among the main hits (Table 1), and it was also reported in small EVs^5^. Interestingly, analysis performed through the String database^25^ indicated three groups of known protein interactions: Flot1-Flot2, Rab7A-DYNLL1, and LCP1-CTTN-ACTG1 (Suppl. Fig. S2C).

**Figure 3:**
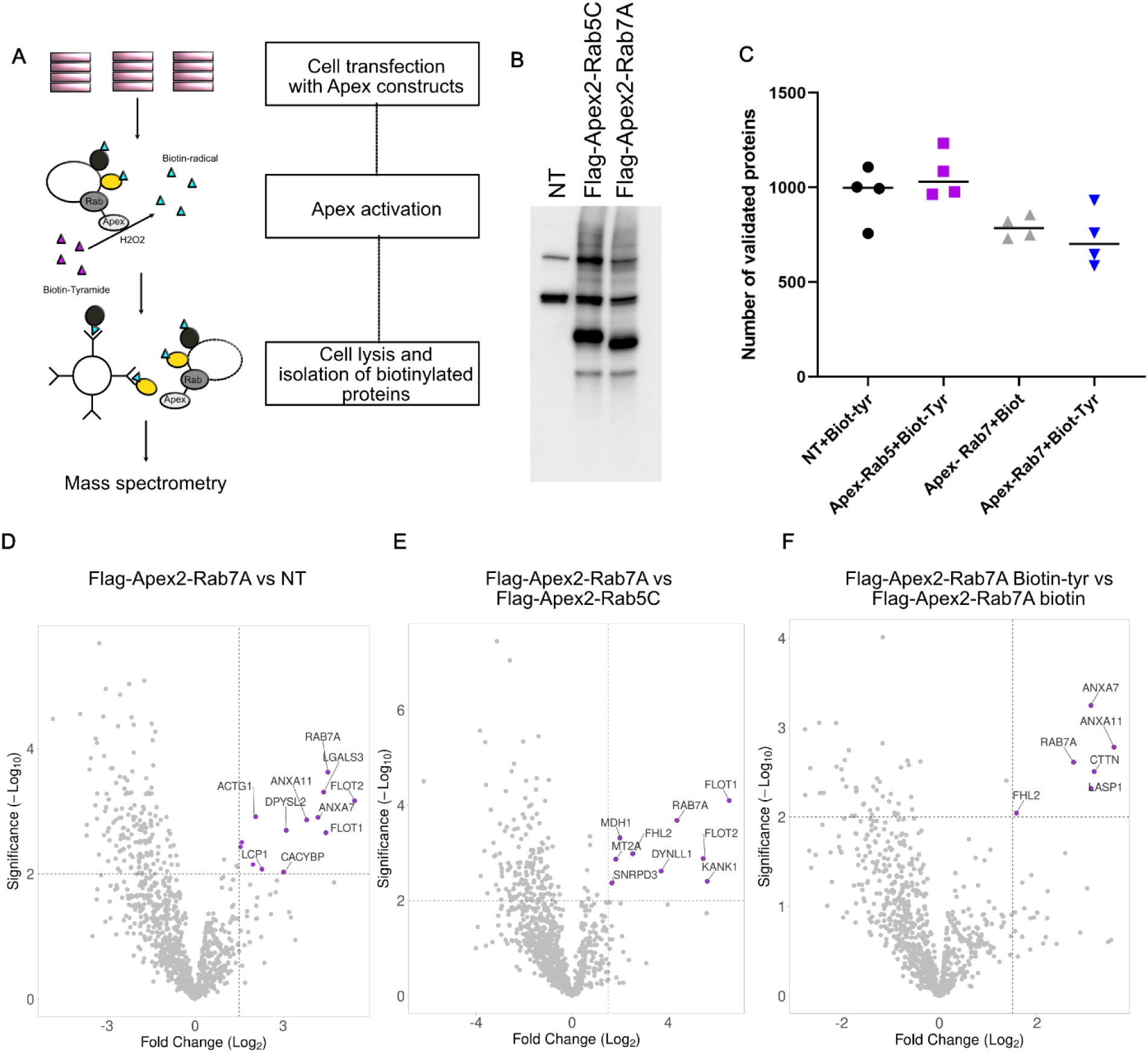
Proteomic study of Rab7A partners identified EV containing proteins. (A) Schematic representing the pipeline used for mass spectrometry. (B) Western blot analysis visualizing the biotinylated proteins after Apex2 activation. SUM159 cells were non-transfected (NT) or transfected with Flag-Apex2-Rab5C and Flag-Apex2-Rab7A. At 24h post-transfection, the Apex2 was activated with hydrogen peroxide in presence of biotin-tyramide. A quantity of 40 µg of proteins was loaded on a gradient gel and a biotin antibody was used to observe the biotinylated partners. (C) Number of proteins validated after LC-MS/MS tandem mass spectrometry. The graph represents the number of proteins identified in quadruplicate in the conditions non-transfected with biotin-tyramide (NT+Biot-tyr), in Flag-Apex2-Rab5C with biotin-tyramide (Apex-Rab5+Biot-Tyr), in Flag-Apex2-Rab7A with biotin (Apex-Rab7+Biot) and in Flag-Apex2-Rab7A with biotin-tyramide (Apex-Rab7+Biot-Tyr). (D) Volcano plot comparing Flag-Apex2-Rab7A with the non-transfected with biotin-tyramide (NT) (E) Volcano plot comparing Flag-Apex2-Rab7A with the Flag-Apex2-Rab5C. (F) Volcano plot comparing Flag-Apex2-Rab7A with biotin-tyramide (Biotin-Tyr) and Flag-Apex2-Rab7A with biotin. Proteins that are selected are named on the volcano plot.

**Table 1.** Main candidates identified by quantified mass spectrometry.

| Conditions | Protein name | Fold change | pvalue | Number of peptides | Protein score |
| --- | --- | --- | --- | --- | --- |
| Rab7A_biotin-tyramide versus NT_biotin-tyramide | Flot2 | 43.1199643 | 0.000678185 | 29 | 1028.181 |
|  | Rab7A | 22.9302609 | 0.000237194 | 29 | 1072.04 |
|  | Flot1 | 21.8772218 | 0.002195434 | 31 | 1227.465 |
|  | LGALS3 | 20.7578727 | 0.000492336 | 7 | 278.469 |
|  | ANXA7 | 18.1733584 | 0.001247898 | 17 | 561.455 |
|  | ANXA11 | 13.9008567 | 0.001368018 | 13 | 437.653 |
|  | DPYSL2 | 8.58915648 | 0.002015316 | 29 | 1170.103 |
|  | ACTG1 | 4.17342829 | 0.001218162 | 40 | 1845.31 |
|  | MT2A | 3.01484182 | 0.00312025 | 1 | 43.531 |
| Rab7A_biotin-tyramide versus Rab7_biotin | ANXA11 | 11.9094155 | 0.001655834 | 13 | 437.653 |
|  | ANXA7 | 8.58178208 | 0.00056556 | 16 | 561.455 |
|  | CTTN | 8.98817056 | 0.003100276 | 24 | 732.684 |
|  | Rab7A | 6.71512177 | 0.002437321 | 23 | 1072.04 |
| Rab7A_biotin-tyramide versus Rab5C_biotin-tyramide | Flot1 | 93.5025764 | 7.97728E-05 | 31 | 1227.465 |
|  | Flot2 | 44.1680546 | 0.001320071 | 29 | 1028.181 |
|  | Rab7A | 20.6466242 | 0.000208224 | 29 | 1072.04 |
|  | DYNLL1 | 13.097523 | 0.002415385 | 6 | 474.372 |

### Rab7A interacts with Flot1

Proteomic studies using chemical crosslinkers and superparamagnetic isolation of lysosomes already identified the presence of Flot1 and Flot2 in endo-lysosomal compartments^26^. Indeed, the role of Flot1 and Flot2 is well defined in endocytosis, and in the recycling of receptors from the endosomes to the plasma membrane^27,28,29^. They act as membrane associated scaffolding proteins that form heterotetramers at the cytosolic face of the cells during endocytosis. However, their role in EV biogenesis or release is still unknown. To our knowledge, Flot1/2 have never been identified as interactors of Rab7A, although they were among our most significant hits in our mass spectrometry analysis (Table 1 and Suppl. Fig. S2C). To determine whether Flot1 and Rab7A can interact (directly or indirectly), we immunoprecipitated Flag-Apex2-Rab7A using an anti-Flag antibody. Through western blot analysis, we noticed a weak interaction between Flot1 and Flag-Apex2-Rab7A (Fig. 4A). This interaction could be transient considering the dynamic switching of small Rab-GTPases from an active to an inactive form through GTP hydrolysis. Thus, we used the membrane-permeable crosslinker dithiobis-(succinymidylpropionate) (DSP) to strengthen the binding between these proteins. We observed that Flot1 bound strongly to Flag-Apex2-Rab7A in the presence of DSP, while Lamp1, another endo-lysosomal protein, was not, supporting the specificity of this interaction. As a control, we showed that the Rab7A effector RILP also interacts with Flag-Apex2-Rab7A in the absence of DSP, and more strongly in its presence, further validating our assay. To monitor the localization of endogenous Flot1 in relation to Flag-Apex2-Rab7A, we performed immunofluorescence assays. Flot1 exhibited a disperse punctate pattern throughout the cytosol with some plasma membrane localization (Fig. 4B-C). A subset of Flot1 colocalized with Rab7A (14.33% +/- 1.21) and CD63 (27.60 +/- 2.08), while significantly weakly colocalizing with the early endosomal marker EEA1 (6.93% +/-0.57). Flot2 localization was also monitored and exhibited a similar profile of colocalization with Rab7A as Flot1 (Suppl. Fig. S3). These findings support our mass spectrometry data (Fig. 3) and reveal that Flot1 is an interacting partner of Rab7A.

**Figure 4:**
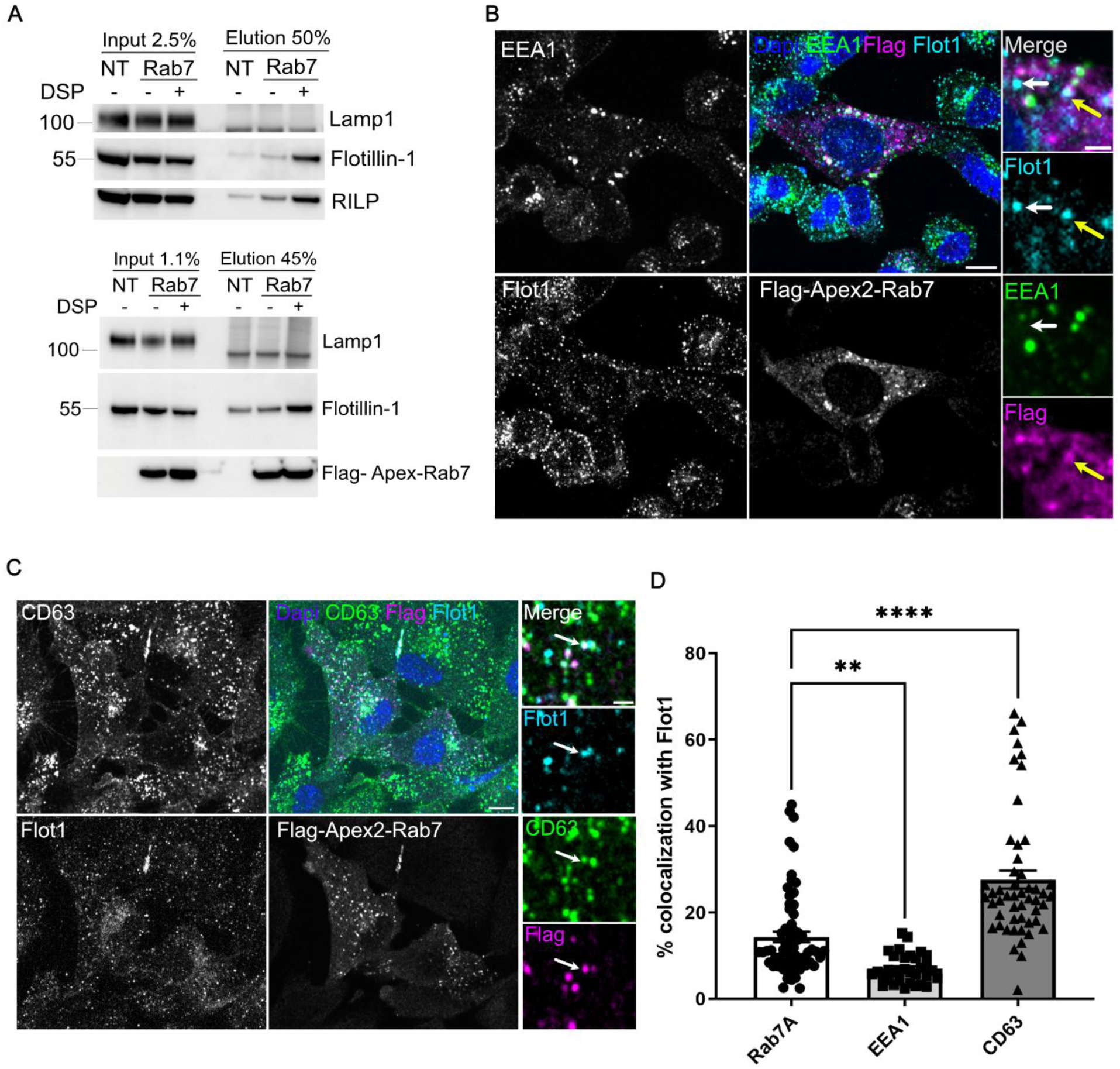
Flotillin1 interacts with Rab7A. (A) Binding assay indicating Flotillin-1 (Flot-1) binding to Rab7A in cross-linking conditions. SUM 159 cells transfected or non-transfected (NT) with Flag-Apex2-Rab7A are placed 2 h in the absence of in the presence of the permeable crosslinker DSP. The cell lysate was collected, and the post-nuclear supernatant was incubated with an agarose resin containing Flag antibodies for immunoprecipitation. The western blot analysis represents the input and eluted fractions. Specific antibodies against Lamp1, Flotillin 1, RILP and Flag-Apex2-Rab7A are used for visualization. The experiment was performed in the represented duplicates. (B) EEA1 does not colocalizes with Flot1. SUM159 cells were transfected with Flag-Apex2-Rab7A. 24h post transfection cells were visualized by confocal microscopy. Representative images from a single z-stack indicated EEA1 (green), Flot1 (cyan) and Rab7A (magenta) using the anti-Flag antibody. White arrows indicate no colocalization between Flot1 and EEA1. Yellow arrows indicate colocalization between Flot1 and Flag-Apex2-Rab7A. Scale bar = 10 µm and scale bar for zoomed regions = 2 µm. (C) CD63 localizes with Flotillin1. SUM159 cells were transfected with Flag-Apex2-Rab7A. At 24 h post-transfection cells were visualized by confocal microscopy. Representative images from a single z-stack indicated CD63 (green), Flot1 (cyan) and Rab7 (magenta) using the anti-Flag antibody. White arrows indicate colocalizations between Flot1 and CD63. Scale bar = 10 µm and scale bar for zoomed regions = 2 µm. (D) The graph represents the percentage of colocalization of Flotillin 1 with CD63, Flag-Apex2-Rab7A, and EEA1. Images from (B-C) were used. Data were plotted as percentage of colocalized volumes +/- s.e.m. Two-way anova test with multiple comparisons were used. **pvalue=0.0040, ****pvalue<0.0001. Each dot represents 30 to 65 field of views from n = 3 independent experiments.

### Rab7A biotinylated partners are identified in EVs

Next, we wanted to evaluate the presence of the Rab7A biotinylated partners within EVs secreted by cells. To detect the presence of biotinylated EVs, the cell culture supernatant of SUM159 cells expressing Flag-Apex2-Rab7A was collected. EVs were isolated using a protocol of differential ultracentrifugation (dUC; see Materials and Methods). Cell debris were excluded in the 2K pellet. The 10K fraction contains large EVs and the 100K fraction small EVs, which includes small microvesicles and exosomes. Following EV isolation, we recovered a higher protein concentration in the 10K and 100K pellets expressing Flag-Apex2-Rab7A compared to the mock-transfected (mock) samples (Fig. 5A). This finding is in accordance with a previous study showing that Rab7A overexpression led to an increase in EV secretion^19^. Focusing on the 100K pellet that likely contains exosomes, we performed nanoparticle tracking analysis (NTA) to characterize the size of the isolated EVs. The size of the mock EVs (126.3 +/- 21.7) was not significantly different from the EVs isolated in the presence of Flag-Apex2-Rab7A (138.8 +/-22.5), (Fig. 5B, and Table 2). This result indicates that Rab7A expression does not significantly affect the size of EVs present in the 100K pellet. Protein characterization of these EVs by western blotting revealed the presence of CD63 (Fig. 5C, D), CD9 (Fig. 5 C, E), and Flot1 (Fig. 5C, F) in the 100K pellet. GP96, an ER associated protein, was undetectable from the 10K and 100K EV pellet samples, indicating that the EV preparation had minimal cell contamination. Interestingly, the expression of Flag-Apex2-Rab7A affected the content of EVs from the 10K pellet by increasing the amount of Flot1 (Fig. 4C, F), and CD9 inside these vesicles (Fig. 4C, E), although the size of these 10K EVs (271.0 +/- 18.0; Table 2) was not significantly different from the mock transfected (241.5 +/- 86.7; Table 2). This data suggests that Rab7A may play a role in the loading of these proteins inside EVs.

**Figure 5:**
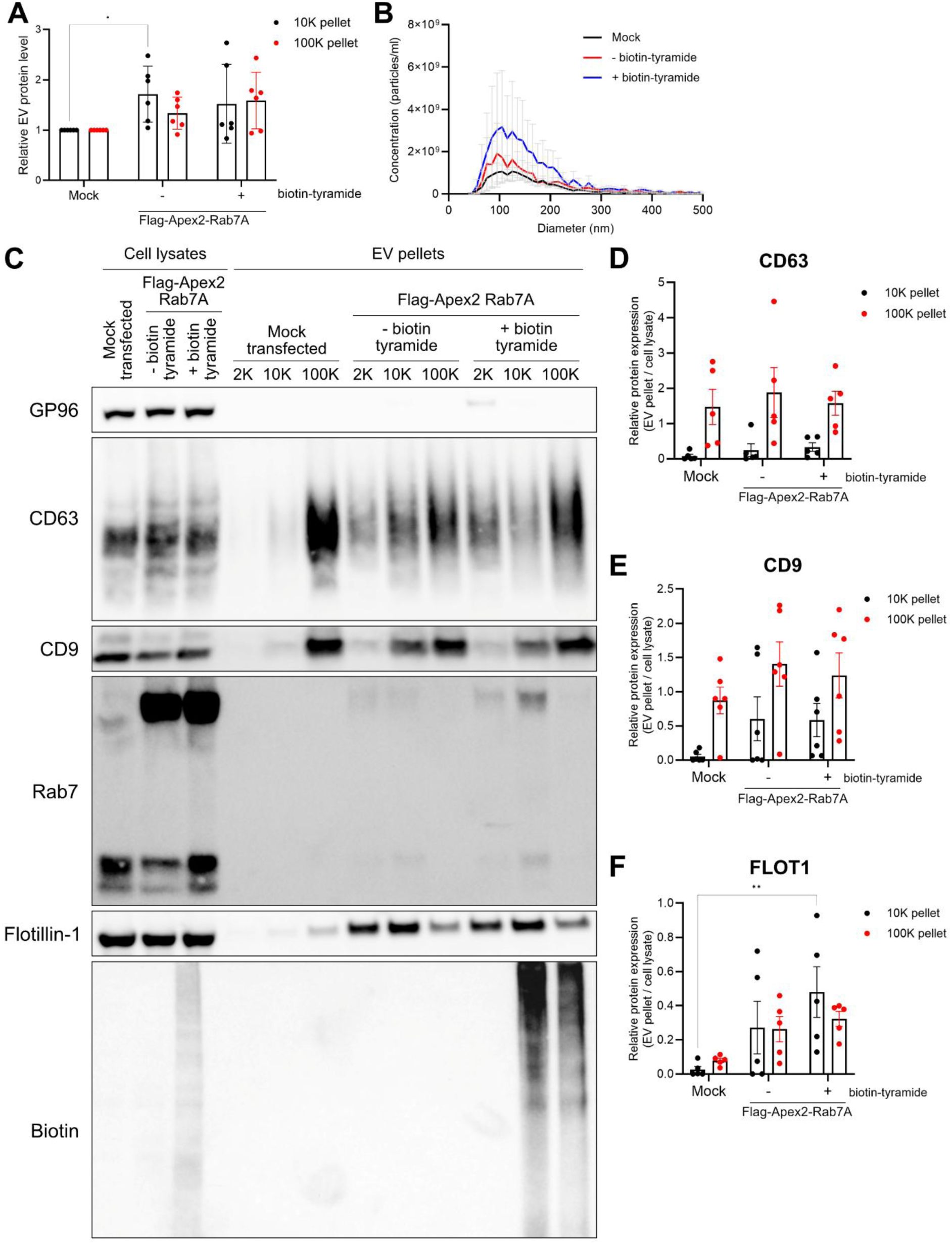
Flag-Apex2-Rab7A biotinylates EVs and modulates EV secretion. (A) Quantification of total protein levels (relative to mock transfected cells) of EV pellets (10K, 100K pellets) isolated from cell culture supernatants via differential ultracentrifugation. Data is plotted as mean +/- s.e.m., n = 5 independent experiments. (B) NTA analysis of 100K pellet isolated from cell culture supernatants of cells either mock-transfected (black line) or expressing FLAG-APEX2-Rab7A WT (red) or DN (blue). Data is plotted as mean +/- s.e.m., n = 3 independent experiments. (C) Western blot analysis of cell lysates and EV pellets (10K and 100K) isolated from mock-transfected cells or expressing FLAG-APEX2-Rab7A WT or DN. (D-F) Quantification of CD63 (D), CD9 (E) and Flotillin-1 (F) (FLOT1) protein expression in EV pellets compared to whole cell lysates. Data is plotted as +/- s.e.m, n = 5 independent experiments. *, p < 0.05 (ordinary two-way ANOVA with a Dunnett’s multiple comparisons test).

**Table 2.** Summary of EV size and concentration from all experiments. Data in the table represent n=3 independent experiments +/-s.e.m

| EV pellet | Condition | Peak (nm) | Median (nm) | Concentration (particles/ml) |
| --- | --- | --- | --- | --- |
| 10K | Mock transfected | 241.5 (± 86.7) | 265.0 (± 79.2) | $1.07 \times 10^9$ (± $4.13 \times 10^8$) |
| | APEX2-Rab7a WT, no biotin | 281.8 (± 44.3) | 325.2 (± 16.0) | $3.25 \times 10^9$ (± $1.95 \times 10^9$) |
| | APEX2-Rab7a WT | 271.0 (± 18.0) | 311.8 (± 19.6) | $2.94 \times 10^9$ (± $2.50 \times 10^9$) |
| | APEX2-Rab7a DN | 254.1 (± 13.2) | 327.5 (± 14.5) | $3.94 \times 10^9$ (± $2.36 \times 10^9$) |
| 100K | Mock transfected | 126.3 (± 22.7) | 144.2 (± 29.3) | $1.47 \times 10^{10}$ (± $4.95 \times 10^9$) |
| | APEX2-Rab7a WT, no biotin | 130.3 (± 21.7) | 155.6 (± 30.9) | $4.51 \times 10^{11}$ (± $7.36 \times 10^{11}$) |
| | APEX2-Rab7a WT | 138.8 (± 22.5) | 162.2 (± 29.5) | $4.05 \times 10^{10}$ (± $5.14 \times 10^{10}$) |
| | APEX2-Rab7a DN | 136.4 (± 21.1) | 161.4 (± 28.0) | $2.47 \times 10^{10}$ (± $2.56 \times 10^{10}$) |

To further characterize the EVs obtained from the Flag-Apex2-Rab7A transfected cells, density gradients were performed on EVs isolated from the 100K pellet and revealed the presence of endogenous Rab7A inside these EVs, supporting the fact that exosomes are included in the 100K pellet as expected (Fig. S4B). In addition, we noticed that expression of Flag-Apex2-Rab7A slightly switched the presence of its proximal partners, Flot1 and Annexin A7, into a less dense fraction (fraction 4), that might correspond to so-called “lipid rafts”. Despite these differences, most of the EV protein markers remained in the same fractions as in the mock sample (fractions 6-10). This observation showed that Rab7A overexpression did not affect the density of these EVs. As shown in Suppl. Fig. S1, we did not observe Flag-Apex2-Rab7A localization at the plasma membrane, which suggest that the presence of the endogenous Rab7A in EVs comes in great part from its loading inside ILVs at the endosomes. Monitoring the biotinylation of Rab7A partners, we were able to identify them in isolated EVs in the 10K and 100K fractions (Fig. 5C). These biotinylated EVs were also found in the fraction containing the EV markers CD63, Flot1, ANXA7 (fraction 6-10), further supporting that they are packaged in EVs of endosomal origin (Fig. S4B). Using the TMR-tyr, we wanted to monitor the presence of fluorescent EVs isolated from the 100K pellet (see Materiel & Methods). We expressed CD63-EGFP, Flot1-EGFP and Flot2-EGFP in SUM159 cells to locate EVs in the extracellular medium. The secreted EVs from the 100K pellet were incubated with SUM159 cells and the presence of fluorescent EVs attached to the cells were monitored by confocal microscopy. We found that TMR-tyr colocalized with CD63-EGFP (Suppl. Fig. S5A), Flot1-EGFP (Suppl. Fig. S5B), and Flot2-EGFP (Suppl. Fig. S5C). Nevertheless, this data illustrates that Rab7A-originating EVs (which are likely exosomes) can readily be labeled in an unpurified mixture of heterogeneous EV subtypes.

## Discussion

EVs are heterogenous in content which is mostly explained by their site of biogenesis. Determining their origin is a challenging task as they contain common markers and are mixed in the extracellular space. Originating from the plasma membrane or endosomes, EVs may have different signaling functions, due to specific proteins at their surface or to the composition of their internal cargoes. Thus, it is crucial to understand the molecular mechanisms that govern their biogenesis and release from cells. To this end, it is essential to develop innovative tools to label specifically EVs at their place of origin, which would improve the identification of a variety of EV subsets and allow the tracking of their release from cells.

In our study, we used a proximity labeling assay to identify EVs of an endosomal origin. We used the small Rab GTPase Rab7A as a bait, considering its MVB localization, to label proteins that would potentially be internalized into ILVs. Thus, to perform the PLA, we engineered a construct in which Rab7A was fused to the Apex2 under the control of a weak promoter (Flag-Apex2-Rab7A). Using quantitative mass spectrometry on biotinylated fractions, we identified several proteins known to be present in EVs. Among them, Flot1 was highly represented. It is described as a scaffold protein commonly found in EVs^17^. Although Flot1 is known to be present in late endosomes, we demonstrated its interaction with Rab7A. Through the characterization of EVs secreted from our cell type, we confirmed the presence of Flot1 and surprisingly identified Rab7A as part of their composition. This observation supported the fact that we isolated bonafide exosomes. More interestingly, EVs isolated from cells expressing Flag-Apex2-Rab7A contained biotinylated proteins, suggesting that Rab7A proximity partners are introduced during ILV formation. By monitoring Flot1, we noticed that the expression of Flag-Apex2-Rab7A increased the secretion of EVs containing Flot1 and increased its loading inside small and large EVs, contrary to CD63. From our study, we demonstrated that our PLA tool Flag-Apex2-Rab7A can specifically label exosomes. In addition, we might have characterized the existence of a new subset of EVs positive for Flot1 and Rab7A that are distinct from CD63 containing EVs.

To use the PLA with Rab proteins, it is important to verify the proper localization of our Rab constructs as well as their specificity of labeling. We monitored Rab5C, Rab6B and Rab7A fused to the Apex2. In the case of Rab7A and Rab5C, we had a double localization of the proteins, in the cytosol and at membrane structures. This double localization was previously known as Rab proteins are in the cytosol as an inactive state bound to GDP and complexed with GDI, while their active state, bound to GTP, is localized at the membrane^30^. Surprisingly, we mostly detected biotinylation at membrane structures which suggest that most of the proximity partners would be identified at vesicular membranes rather than the cytosol. Using PLA and the fluorescent molecule TMR-tyramide, we were able to induce the labeling of Rab7A proximity partners in a timely manner and monitor by live imaging their trafficking within Rab7A positive vesicles. Using TMR-tyramide, we could track the mobility of a pool of proximity partners, but we do not have so far, the resolution to discriminate Rab7A intracellular partners from the ones that would be part of EV secretion. Through the monitoring of EVs secreted from cells expressing Flag-Apex2-Rab7A and incubated with TMR-Tyramide, we could identify few fluorescent particles that are double positives for TMR and the EV markers CD63-EGFP, EGFP-Flot1 or EGFP-Flot2. The limitation of this assay relies on the quantity of fluorescent partners that would be packaged inside a single EV as well as the number of fluorescent molecules needed to be visualized by confocal microscopy. Using confocal imaging and fluorescent correlation spectroscopy on single EV containing CD63-EGFP, it was suggested that a range between 5 to 10 molecules of EGFP would be needed to observe a single fluorescent EV particle^31^.

To identify Rab7A proximity partners, we used quantitative proteomics. The PLA is often combined with proteomics at the cellular or tissue level to decipher proteins acting in specific pathways^22,32^. However, quantitative proteomics remains a difficult task as we can identify many nonspecific proteins. To improve the selection of relevant candidates, we used two nonspecific controls (non-transfected cells with biotin-tyramide, Rab7A transfected cells with biotin) and Flag-Apex2-Rab5C as a control of specificity. Despite these controls, we identified many proteins in each condition (<700 proteins). The subtraction of our controls from the sample of study (Flag-Apex2-Rab7A in presence of biotin-tyramide) allowed us to identify relevant partners such as the Dynein light chain 1. Dynein is known to be part of the Dynein-dynactin complex interacting with Rab7A and RILP for the retrograde transport of vesicles from late endosomes to lysosomes^33,34^. This result highlighted the specificity of our assay. However, we did not identify any other Rab7A effectors such as RILP potentially due to the absence of tyrosine residues at proximity of the Apex2 which is a limitation to consider. Another proteomic study performed on Apex2-Rab21 used Apex2-Rab7A as a control of specificity. They identified proximity partners from the retrograde transport and few known Rab7A effectors such as Vps35 and the GAP TBC1D5^35^. Compared to our study, these differences could be explained by the fact that Rab7A was overexpressed with the strong CMV promoter, in addition they are using a different cell line, where Rab7A expression could potentially have a stronger role in the retrograde transport. In our SUM159 cell line, most of the identified candidates were classified by the Gene Ontology-term analysis as being present in extracellular vesicles. The annexins A7 and A11 were found in small EVs although their role is still unknown^5^. Annexins are calcium sensitive proteins mostly described in membrane repair processes. They bind to phospholipids and were visualized at the plasma membrane or in intracellular vesicles^36^. Annexin A7 was recently demonstrated as acting in a lysosomal repair process independently of ESCRT III^37^. Annexin A11 was identified as a molecular tether binding RNA granule to lysosomes. It is known that exosome release depends on calcium level^38,39^. Another annexin such as annexin A6 was suggested to tether MVBs to the plasma membrane in a calcium dependent manner^39^. Annexin A6 was also demonstrated to recruit TBC1D15, the GAP for Rab7A^40^. The binding of Rab7A to Annexin A7 and A11 could have a regulatory function and their link with exosome biogenesis or release needs to be investigated^41^.

We focused our study on Flot1 and Flot2 which are found in EVs and are suggested to form ILVs in an ESCRT-independent manner^15^. The role of Rab7A in the biogenesis of ILVs is unclear. It was demonstrated that Rab7A depletion had no impact on ILVs size, and is not needed for the formation of ILVs^42^. Moreover, its role on EV secretion is not elucidated as Rab7A deficiency could either decrease or increase the release of total secreted EVs^20,43^. Another report suggested that Rab7A acted on EVs secretion upstream of the MVB-PM fusion events. In this study, ER-membrane contact sites would favor the maturation of late endosomes, switching Rab7A to Arl8B to direct the MVB transport toward the PM for EV secretion ^19^. Although, we do not study these steps, our data supports that Rab7A acts upstream of MVB-PM fusion. In our study, we were not able to observe ILVs by transmission electron microscopy, although we did notice by NTA that the secreted EVs preserved their regular size compared to mock transfected cells. In addition, we observed that Rab7A overexpression led to an increase in total protein content and EV particle number. More interestingly, among the EV markers monitored, this increase only concerns Flot1 and CD9 but not CD63. This specific effect of Rab7A overexpression is surprising as in other studies deficiency or overexpression of Rab7A impact most of the common EV markers^19,20^. By density gradient, we noticed the presence of a lighter fraction containing Flot1, AnxA7 but not CD63. Flot1 and Annexins are known to be present in microdomains where they could act as platforms to recruit specific proteins^44,45^. This shift in vesicle density could suggest the presence of these proteins inside specific lipid microdomains which are triggered by the expression of Rab7A. One hypothesis would be that Rab7A could be recruited by these proteins and increase the formation of these microdomains. Microdomains are usually needed to sort cargoes. Considering the suggested role of Flot1 and Flot2 in membrane curvature^46^ and its ability to interact with their cargoes^26^, we suggest that Rab7A combined with Flot1 could allow the recruitment of specific cargoes which will be introduced inside ILVs. The increase in Rab7A expression could increase the probability of formation of these microdomains, recruiting specific proteins which will be loaded in large and small vesicles formed at the MVBs. This would explain the presence of Flot1 and biotinylated proteins inside large and small EVs. It was demonstrated that lysosome inhibition could also induce the secretion of large EVs^43^. Thus, we cannot exclude the fact that Rab7A overexpression could change the balance between its role at the MVBs *versus* its function in protein trafficking from the late endosome to the lysosome.

Collectively, our study demonstrated that through Rab7A, we could label proteins that are introduced inside ILVs, supporting the fact that Rab7A protein partners could be loaded at the late endosome stage. By discovering Rab7A protein partners, we opened new insights in proteins that might have a role in cargoes selection in the ILVs. Developing Flag-Apex2-Rab7A as a tool, we demonstrated that we could label EVs and potentially track them intracellularly. More importantly, we might have uncovered a new mechanism of ILVs loading involving Rab7A and Flot1 in the formation of specific EV subtypes.

## Methods

### Cell culture

SUM159 cells were maintained in Dulbecco’s modified Eagle’s medium (DMEM)-F12, Glutamax (Gibco), 5% fetal bovine serum (FBS, Sigma-Aldrich), 1% penicillin/streptomycin (Gibco), Insulin (5ug/ml, Sigma-Aldrich), hydrocortisone (1ug/ml, Sigma-Aldrich), HEPES pH 7.0 (10mM, Gibco). Cells were maintained at 37°C, 5% CO_2_.

### Cloning

Flag-Apex-Rab5C (Addgene plasmid # 166404), Flag-Apex2-Rab6B (Addgene plasmid # 166398), Flag-Apex2-Rab7A (Addgene plasmid #135652), EGFP-Rab7A (Addgene plasmid # 226184) cDNA constructs were cloned into pBS vectors under the control of the weak promoter L30 and were generated by the Montpellier Genomic Collection (MGC).

The ALFA-Apex2-Rab7A (Addgene plasmid #224892) construct was generated from Flag-Apex2-Rab7A using the Q5 Site-Directed mutagenesis kit to replace the Flag tag. Primers used were: forward primer 5’ GAGACGCCGCTTAACTGAACCTGGAAAGTCTTACCCAACTG3, reverse primer 5’ AGTTCCTCTTCCAAACGTGATGGCATGGTGGCGGATCCTTA 3’.

For live imaging studies, SUM159 cells stably expressed the nanobody miRFP680. They were generated after lentiviral transduction with the plasmid lenti NbALFA-miRFP680 puro (Addgene plasmid# 193972). Cells were selected after 10µg/ml puromycine selection. Other versions of these plasmids are available for the community: Lenti NbALFA-miRFP680 hygro (Addgene plasmids #193969), Lenti NbALFA-mNeonGreen puro (Addgene plasmids#193973) and Lenti NbALFA-mNeonGreen hygro (Addgene plasmids #193974).

Flot1-EGFP, Flot2-EGFP and CD63-EGFP were a kind gift from Stephane Bodin at CRBM Montpellier.

### Antibodies and treatments

For immunofluorescence assay, the primary antibodies used were Rabbit anti-Flot1 (1:100; Cell Signaling Technology), Rabbit anti-Flot2 (1:100, Novus Biologicals), Goat anti-Flag (1:500, Novus biologicals), mouse anti-CD63 (1:1000, ThermoFisher Scientific), mouse anti-EEA1 (1:1000, BD Biosciences), mouse anti-Lamp1 (1:1000,BD Biosciences), sheep anti-TGN46 (1:1000, Novus biologicals), Rabbit anti-Flag (1:500,), mouse anti-Biotin (1:500, Novus Biologicals). Secondary antibodies used were Donkey Alexa Fluor plus 647 anti-rabbit (1:1000, ThermoFisher Scientific), Donkey Alexa Fluor plus 488 anti-mouse (1:1000, ThermoFisher Scientific), Donkey Alexa Fluor 561 anti-sheep (1:1000, ThermoFisher Scientific), DAPI (4′,6-diamidino-2-phenylindole) (1:1000; Sigma-Aldrich) for nucleus staining.

For immunoblotting, the primary antibodies used were rabbit anti-Rab7A (1:1000, Cell signaling technology), rabbit anti-Flot1 (1:1000, Cell signaling technology), mouse anti-Lamp 1 (1:1000, BD Biosciences), rabbit anti-Flag (1:1000,), mouse CD63 (1:1000, ThermoFisher Scientific), mouse CD9 (1:1000, Merck Millipore), mouse GP96(1:1000, Enzo Life Sciences), mouse anti-Biotin (1:500, Novus Biologicals), rabbit anti-Annexin A7 (1:1000, Proteintech) rabbit-Anti RILP (1:500, Novus Biologicals), mouse anti-GAPDH (1:1000, Genetex). The secondary antibodies used are donkey anti-mouse HRP (1:5000, ThermoFisher Scientific), donkey anti-rabbit HRP (1:5000, ThermoFisher Scientific).

For proximity labeling assay, hydrogen peroxide (500µM or 2mM, Sigma-Aldrich), Biotin-tyramide (500µM, Cliniscience), Trolox (10mM, Sigma-Aldrich), Sodium ascorbate (10mM, Sigma-Aldrich), TMR-tyramide plus (1:50, Akoya Biosciences).

For immunoaffinity assay and crosslinking, anti-Flag M2 agarose beads (Sigma-Aldrich), DSP (Pierce), (150ng/ml), Flag peptide (Sigma Aldrich).

For transmission electron microscopy, SIGMAFAST 3,3’-Diaminobenzidine tablets (DAB) (Sigma-Aldrich).

### Transmission electron microscopy and DAB

SUM159 cells were placed in transwells for 24-well plates at a density of 300,000 cells per well. The next day, they were transfected with Flag-Apex2-Rab7A with Lipofectamine 3000 according to the manufacturer’s protocol. At 24h post-transfection, the medium was removed, and cells were fixed 1 h with 4% paraformaldehyde and put on ice for 5min. Cells were rinsed 5min with 20 mM of cold glycine diluted in PBS. Cells were washed five-time 5min in cold 100 mM sodium cacodylate pH 7.4 containing 2 mM calcium chloride. Cells were incubated 15min with cold Sigma Fast DAB with addition of 0.03% of hydrogen peroxide. The reaction was stopped with 100 mM of cold sodium cacodylate pH 7.4 containing 2mM calcium chloride. Cells were washed five-time 5min in cold 100 mM sodium cacodylate pH 7.4 containing 2 mM calcium chloride. Cells were immersed in a solution of 2.5% glutaraldehyde in PHEM buffer (1X, pH 7.4) overnight at 4°C. They were then rinsed in PHEM buffer and post-fixed in a 0.5% osmic acid + 0.8% potassium Hexacyanoferrate trihydrate for 2h at dark and room temperature. After two rinses in PHEM buffer, the cells were dehydrated in a graded series of ethanol solutions (30-100%). The cells were embedded in EmBed 812 using an Automated Microwave Tissue Processor for Electronic Microscopy, Leica EM AMW. Thin sections (70 nm; Leica-Reichert Ultracut E) were collected at different levels of each block. These sections were counterstained with uranyl acetate 1.5% in 70% Ethanol and lead citrate and observed using a Tecnai F20 transmission electron microscope at 120KV in the Institute of Neurosciences of Montpellier (INM): Electronic Microscopy facilities, INSERM U1298, University Montpellier, Montpellier France.

### Mass spectrometry

#### Sample preparation and proximity ligation assay

SUM159 cells were plated at a density of 1×10^6^ cells in four dishes of 10 cm for each condition. The next day, they were transfected with JetPrime (Polyplus) according to the manufacturer’s instructions with Flag-Apex-Rab7A and Flag-Apex-Rab5C constructs.

At 24 h post-transfection, the cells were washed two times with PBS 1x then incubated 30 min with 500 µM of Biotin-tyramide diluted in complete medium. H_2_O_2_ was added at 500 µM for 10 min, and cells were lysed in RIPA buffer containing 5 mM Trolox, 10 mM sodium ascorbate (Sigma-Aldrich) and protease inhibitors (Promega). The samples were maintained on ice for 10min then they were span 10 min at 10,000 x g at 4°C. The supernatants were collected and incubated overnight with streptavidin agarose resin (Pierce). The following day, the beads were washed three time, and the biotinylated proteins were eluted with Laemmli buffer (10 mM Tris pH 6.8,1 mM EDTA, 5 % ß-mercaptoethanol, 5 % SDS, 10 % glycerol).

### Quantitative Proteomics

Protein concentration of all samples was determined using the RC-DC protein assay (Bio-Rad) according to the manufacturer’s instructions. Two µg of each protein extract were heated at 95°C for 5 min and concentrated on a stacking gel band. Gel bands were cut, washed, reduced with 10 mM dithiothreitol and alkylated using 55 mM iodoacetamide prior overnight trypsin digestion (Promega). The generated peptides were extracted, evaporated under vacuum and resuspended in 10 µL of H20 and 0.1% formic acid before nanoLC-MS/MS analysis.

Extracted peptides were analysed by nanoLC-MS/MS on a nanoUPLC system (nanoAcquityUPLC, Waters) coupled to a quadrupole-Orbitrap mass spectrometer (Q-Exactive HF-X, ThermoFisher Scientific). The solvent system consisted of 0.1% formic acid in water (solvent A) and 0.1% formic acid in acetonitrile (solvent B). Samples (800ng) were loaded into a Symmetry C18 precolumn (0.18 x 20 mm, 5 μm particle size; Waters) over 3 min in 1% solvent B at a flow rate of 5 μL/min followed by reverse-phase separation (ACQUITY UPLC BEH130 C18, 200 mm x 75 μm id, 1.7 μm particle size; Waters) using a 60 minutes linear gradient ranging from 8% to 35% of solvent B at a flow rate of 400 nL/min. A Top 10 method was used with automatic switching between MS and MS/MS modes to acquire high resolution MS/MS spectra. To minimize carry-over, two solvent blank injection was performed after each sample.

NanoLC-MS/MS data was interpreted to do label-free extracted ion chromatogram-based differential analysis. Searches were done using Mascot software (version 2.6.2, MatrixScience) against a *Homo Sapiens* protein sequences database downloaded from SwissProt (03-03-2021; 20.352 sequences, Taxonomy ID 9606) to which common contaminants and decoy sequences were added. Spectra were searched with a mass tolerance of 15 ppm in MS mode and 0.07 Da in MS/MS mode. One trypsin missed cleavage was tolerated. Carbamidomethylation of cysteine residues was set as a fixed modification. Oxidation of methionine residues and acetylation of proteins’ n-termini were set as variable modifications. Identification results were imported into Proline software (version 2.2.0)^47^ and validated. The maximum false discovery rate (FDR) was set at 1% at PSM and protein levels with the use of a decoy strategy. Peptides abundances were extracted without cross assignment between all samples. Protein abundances were computed using the best ion of the unique peptide abundances normalized at the peptide level using the median. To be considered, proteins must be identified in at least three out of the four replicates in at least one condition. The imputation of the missing values and differential data analysis were performed using the open-source ProStaR software (version 1.22.6)^48^. Imputation of missing values was done using the approximation of the lower limit of quantification by the 2.5% lower quantile of each replicate intensity distribution (“det quantile”). A Student t-test was applied on the dataset to perform differential analysis. The adaptive Benjamini-Hochberg procedure was applied to adjust p-values and FDR. The mass spectrometry proteomics data have been deposited to the ProteomeXchange Consortium via the PRIDE partner repository^49^ with the dataset identifier (PXD053702).

### Immunofluorescence assay

SUM159 cells were plated at 100, 000 cells per wells on coverslips of 12 mm diameter. The next day, they were transfected with Flag-Apex2-Rab7A, Flag-Apex2-Rab6B or Flag-Apex2-Rab5C using lipofectamine 3000 according to manufacturer’s protocol. At 24h post-transfection, they were washed one time with PBS and fixed 15 min at room temperature with paraformaldehyde 4%. The cells were washed three times with PBS and incubated with the primary antibodies for 1 h. The cells are washed three times with PBS and incubated with the secondary antibodies for 2h. Cells were washed three times with PBS and mounted with Mowiol 4-88 (Sigma Aldrich). For proximity labeling assay, before fixation, cells were incubated 30min with biotin-tyramide and the Apex2 is activated with 500 µM H2O2 for 10 min.

Cells were visualized with Dragonfly spinning disk confocal microscope. Images were acquired with dragonfly Andor software. The 60X oil objective Apo lambda 1.4 NA DT 0.13 mm and EMCCD iXon888 Life Andor were used.

### Immunoprecipitation assay and crosslinking assay

SUM159 cells were seeded in two 100 mm cell culture dishes at a density of 1×10^6^ cells. The next day, cells were transfected with Flag-Apex2-Rab7A with lipofectamine 3000 according to the manufacturer’s protocol. The cell medium was washed two times with PBS. DSP at 2 mM was mixed in cold PBS and the crosslinker was incubated 2h on ice. No DSP was added in the non-transfected. Cells were washed with medium to inactivate the crosslinker and incubated 15min on ice. Lysis was performed with RIPA buffer and protease inhibitor. Lysate was span at 10, 000 x g for 20 min and incubated overnight at 4°C with anti-Flag M2 beads. The next day, the beads were washed three times with cold PBS. Samples were eluted with Laemmli buffer (ThermoFisher Scientific) in presence of 50 mM DTT. Samples were run on a SDS-PAGE 4-12% Bolt Bis-Tris gradient gel (ThermoFisher Scientific) and transferred onto a nitrocellulose membranes (Bio-Rad).

### EV isolation by differential ultracentrifugation

Cells were seeded in 3 x 150 mm cell culture dishes at a density of 1 x 10^7^ per dish. The following day, cells were transfected with plasmids encoding Flag-Apex2-Rab7A constructs, depending on the experiment. At 16-24 h post-transfection, cells were incubated with media containing biotin-tyramide (500 µM) for 30 min at 37°C, then treated with H_2_O_2_ for 10 min at 37°C to activate the Apex2 enzymatic activity. Cells were then rinsed twice with PBS then incubated overnight in serum-free media. The following day, cell culture medium was collected into 50 ml falcon tubes, and cells were trypsinized and counted. The medium was centrifuged at 300 x *g* for 5 min to remove floating cells. Supernatant was transferred into fresh 50 ml falcon tubes and serial centrifugations of the supernatant at 2000 x *g* (2K), 10,000 x *g* (10K) and 100,000 x *g* (100K) at 4°C was performed. The 2K centrifugation was performed for 20 min using an Eppendorf Centrifuge 5810, and the pellet was discarded. The 10K centrifugation was performed for 40 min, the pellet was washed in 50 ml PBS and recentrifuged for 40 min at 10,000 x *g* in a Fiberlite F15-8 x 50cy Fixed Angle Rotor (ThermoFisher Scientific). The 100K centrifugation was performed for 90 min in a SW32Ti rotor (Beckmann Coulter). The pellet was resuspended in 5 ml of PBS and centrifuged again at the same speed in a SW55Ti rotor (Beckmann Coulter) for 90 min. The 10K and 100K pellets were then resuspended in 100 µl PBS.

For the harvest of fluorescent EVs, cells were seeded in 100 mm cell culture dishes, and were transfected with the relevant plasmids. The following day after transfection, cells were incubated with normal growth media containing TMR-tyramide (1:100, Akoya Biosciences) for 30 min at 37°C. Cells were then treated with 500 µM H_2_O_2_ to activate Apex2 enzymatic activity for 10 min at 37°C. Cells were then washed twice with PBS then incubated overnight in serum-free media. After 16-24 h, cell culture supernatants were collected into tubes and EVs were isolated using the differential ultracentrifugation protocol as previously described. The 100K pellet was added to SUM159 cells seeded into an 8-well imaging chamber (Ibidi) and incubated for 3 h at 37°C. The chamber was then mounted onto a spinning disk confocal microscope to image EV attachment and internalization.

For EV western blots, 5 µg of each EV pellet were run alongside 20 µg of whole cell lysates on a 4-12% Bolt Bis-Tris gradient gel (ThermoFisher Scientific) at 110 V for 1.5 h. Proteins were then transferred onto nitrocellulose membranes using the Trans-Blot Turbo Transfer System (Bio-Rad) using the “MIXED MW” protocol. Membranes were blocked for 30 min at room temperature using 5% (vol/vol) non-fat dry milk in TBS-Tween (TBS-T), 0.05% (v/v) as a blocking buffer. Membranes were probed overnight using primary antibodies diluted in blocking buffer at 4°C on a rocker. The following day, membranes were rinsed three times for 10 min using TBS-T then probed with secondary antibodies conjugated with horseradish peroxidase (HRP) for 1 h at room temperature. Membranes were rinsed again three times for 10 min before incubating with ECL Clarity substrate mix (Bio-Rad) for 5 min at room temperature. Blots were revealed using the ChemiDoc imaging system (Bio-Rad).

### Nanoparticle tracking analysis (NTA)

NTA was performed using ZetaView x30 TWIN (Particle Matrix) equipped with 488 and 640 nm lasers, at 10x magnification, with software version 1.3.8.2. The instrument settings were 25°C, sensitivity of 80, and shutter of 100. Measurements were done at 11 different positions, and frame rate of 30 frames per second.

### Sucrose density gradient

EVs were collected according to the previous protocol. A continuous gradient was prepared with two sucrose solutions (0.4 M and 2 M) diluted in 150 mM NaCl, 50 mM Tris-HCl pH 7.4. The 0.4 M sucrose was layered on top of the 2 M sucrose and the tubes were left overnight horizontally at 4°C to diffuse and form the gradient. The next day, 1 mL of isolated EV resuspended in PBS, were placed on top of the gradient. The tubes were span at 100,000 x *g* for 16h in a SW55Ti rotor (Beckman Coulter). 200 ul of samples were collected and precipitated with trichloroacetic acid (TCA). For the precipitation, 5ul of cold 0.5% (w/v) sodium deoxycholate were added to 200 µl of samples, and 200 µl of cold 10% TCA was added. Samples were vortexed and left on ice for 20 min. Samples were span at 14, 000 x g for 20 min at 4°C. The supernatant was discarded, and the pellet was resuspended in Laemmli buffer for SDS-PAGE analysis.

### Live imaging

Cells were seeded onto 8-well chambered polymer coverslips (Ibidi) and transfected using Lipofectamine 3000 (ThermoFisher Scientific) according to the manufacturers’ instructions. Cells were incubated with TMR-Tyramide (Akoya Biosciences) for 30 min at 37°C, then were treated with 500 µM H2O2 for 10 min at 37°C to activate the Apex2 enzymatic activity. Cells were then washed with media three times for 5 mins to remove any unbound TMR. Live imaging was performed using an Eclipse Ti2 inverted microscope (Nikon) coupled to the Andor Dragonfly spinning disk system (Oxford Instruments), an EMCCD Andor camera (Oxford Instruments), and a 100X APO (1.45 NA) oil objective lens controlled by Andor Fusion software (Oxford Instruments).

### Image analysis

Image processing was performed with Fiji software. Colocalization analyses were performed with Imaris software v9.7 (Bitplane, Oxford Instruments). Surface rendering analysis were performed to select the Flag-Apex2-Rab7A positive cells and used as a mask to exclude non-positive cells. Flag-Apex2-Rab7A positive vesicles were selected through surface rendering to only consider the cytosolic fraction. Colocalizations were performed using an identical threshold for each staining for different images (EEA1, Flot1, Flot2, CD63). This analysis was repeated for n= 2-3 independent experiments. Statistical analyses were performed using Prism version 10.1 (GraphPad). Volcano plots were performed using the software VolcaNoseR^50^. Gene ontology analysis was performed using g: Profiler (version e111_eg58_p18_f463989d)^24^. String analysis were performed using curated database, experiment and co-expression protein data bases ^25^.

## Data availability

All data needed are available in the paper or the supplementary materials. Newly generated plasmids are available through the Addgene platform.

## Acknowledgments

We acknowledge the imaging facility MRI, member of the national infrastructure France-BioImaging (https://ror.org/01y7vt929) supported by the French National Research Agency (ANR-24-INBS-0005 FBI BIOGEN). We acknowledge the Montpellier Genomic Collection (MGC) facility at IGMM member of BioCampus for designing the Apex2-Rab constructs.

## Author contributions

YB, MSD, RG conceived the experiments. MSD, RG, and YB wrote the manuscript. YB, MSD performed the experiments and microscopy analysis. MSD, FD, CC, AH analyzed the proteomics data. Chantal Cazevielle prepared the electron microscopy samples. FD, AH, CC performed the quantitative mass spectrometry analysis. VL generated the nanobody plasmids. MSD wrote the first draft of the manuscript. RG, YB, MSD edited the manuscript. All authors commented on the manuscript.

---

**Figure S1:**
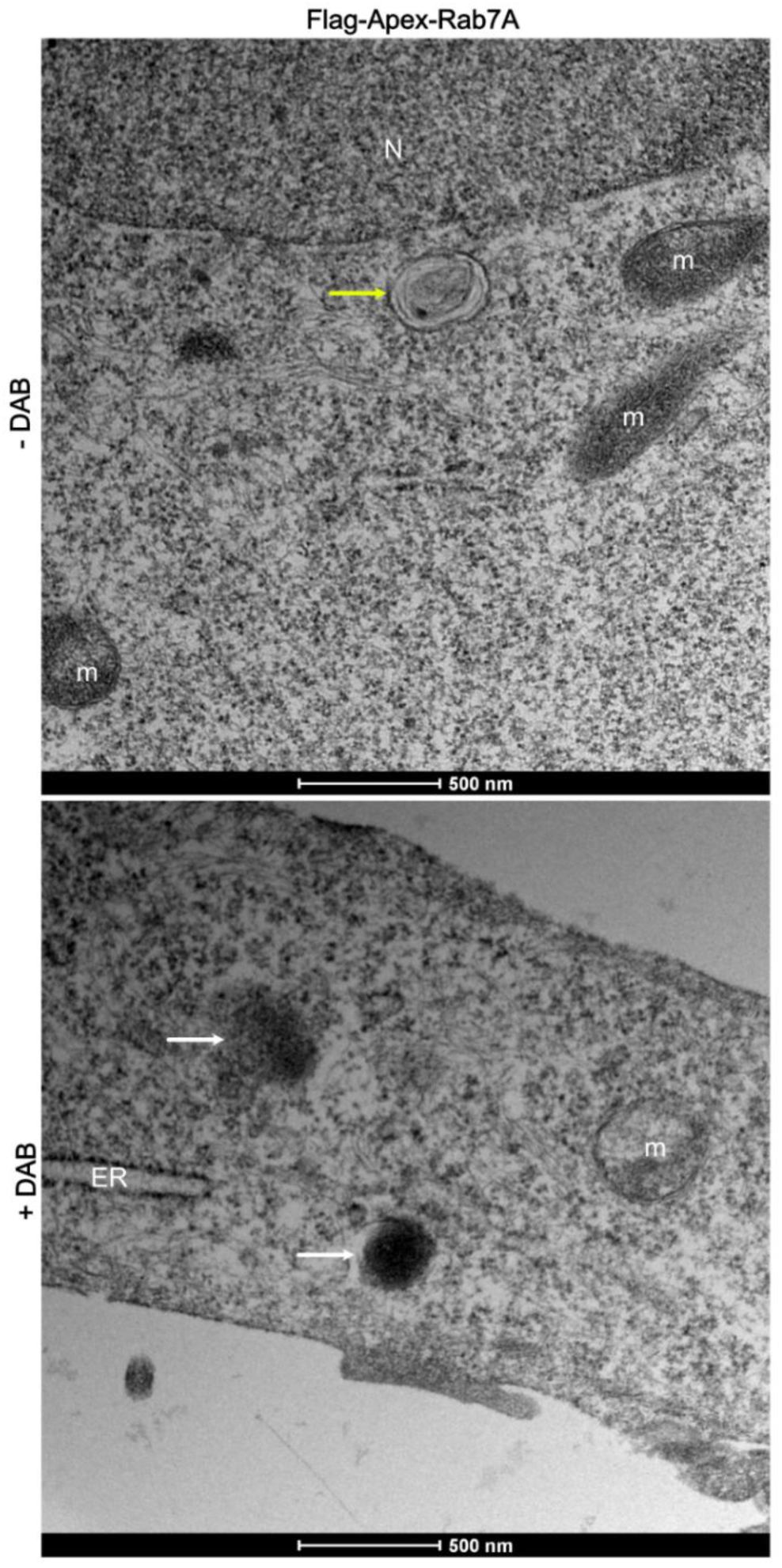
Flag-Apex2-Rab7A localization in cells. SUM159 cells were transfected with Flag-Apex2-Rab7A stained with or without DAB at 24 h post-transfection. Vesicles dense to electrons represented DAB deposits at the localization of Flag-Apex2-Rab7A (white arrow). Vesicles without DAB (yellow arrow), m: mitochondria, N: nucleus, ER: endoplasmic reticulum. Scale bars = 500 nm.

**Figure S2:**
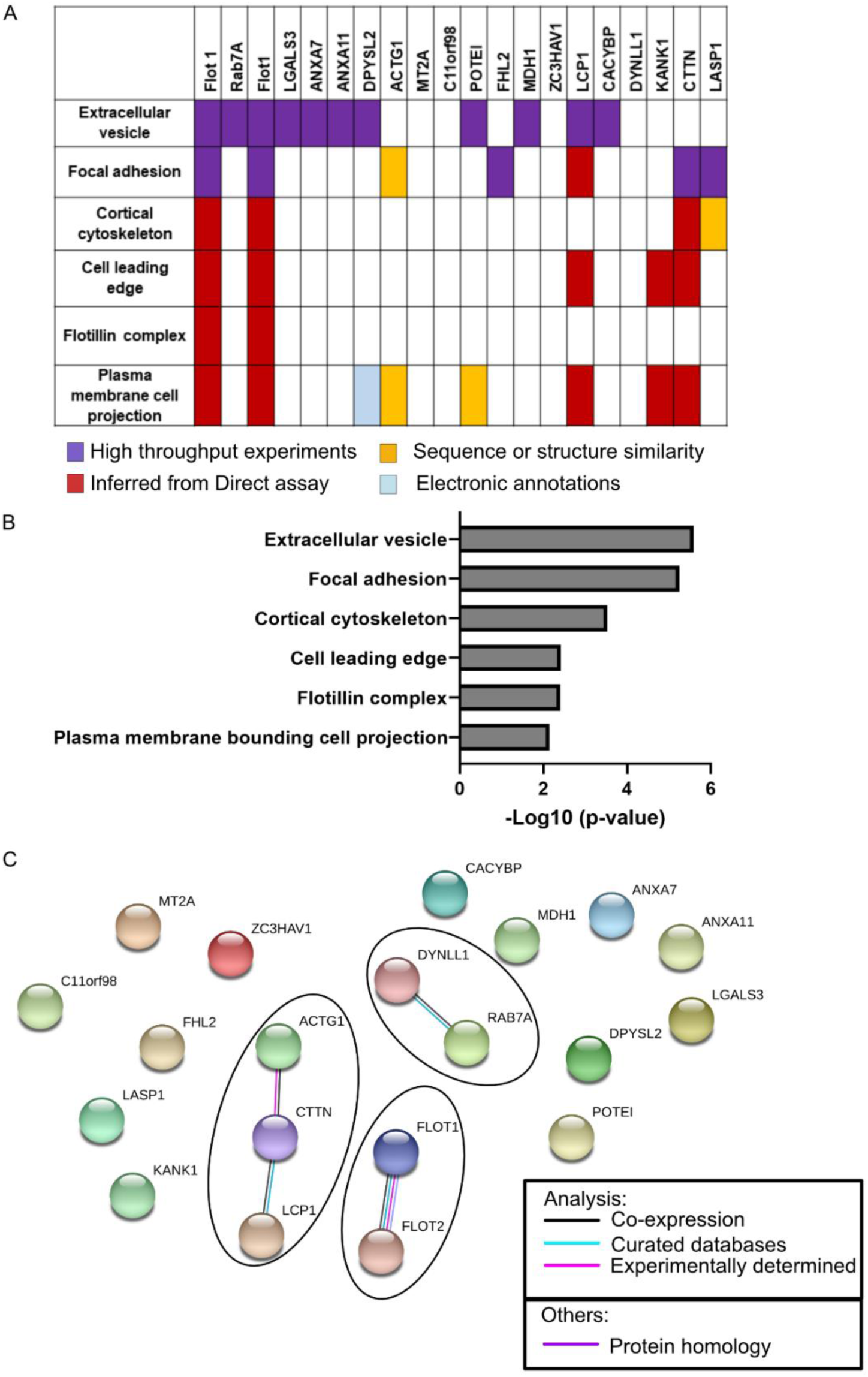
Classification of the Rab7A interacting proteins using cellular components. (A) Gene ontology with the software g:Profiler representing the cellular components. Selected hits were classified according to the different databases: high throughput experiments, inferred from direct assay, sequence or structure similarity or electronic annotations. (B) Significance of the classification in (A) was represented as a bar graph with p-values in -log_10_. (C) String databases identified the link between the different candidates selected in the mass spectrometry analysis. Data from medium confidence were selected, color bonds are representing the different interactions: co-expression, curated databases, experimentally determined interactions, or protein homologies.

**Figure S3:**
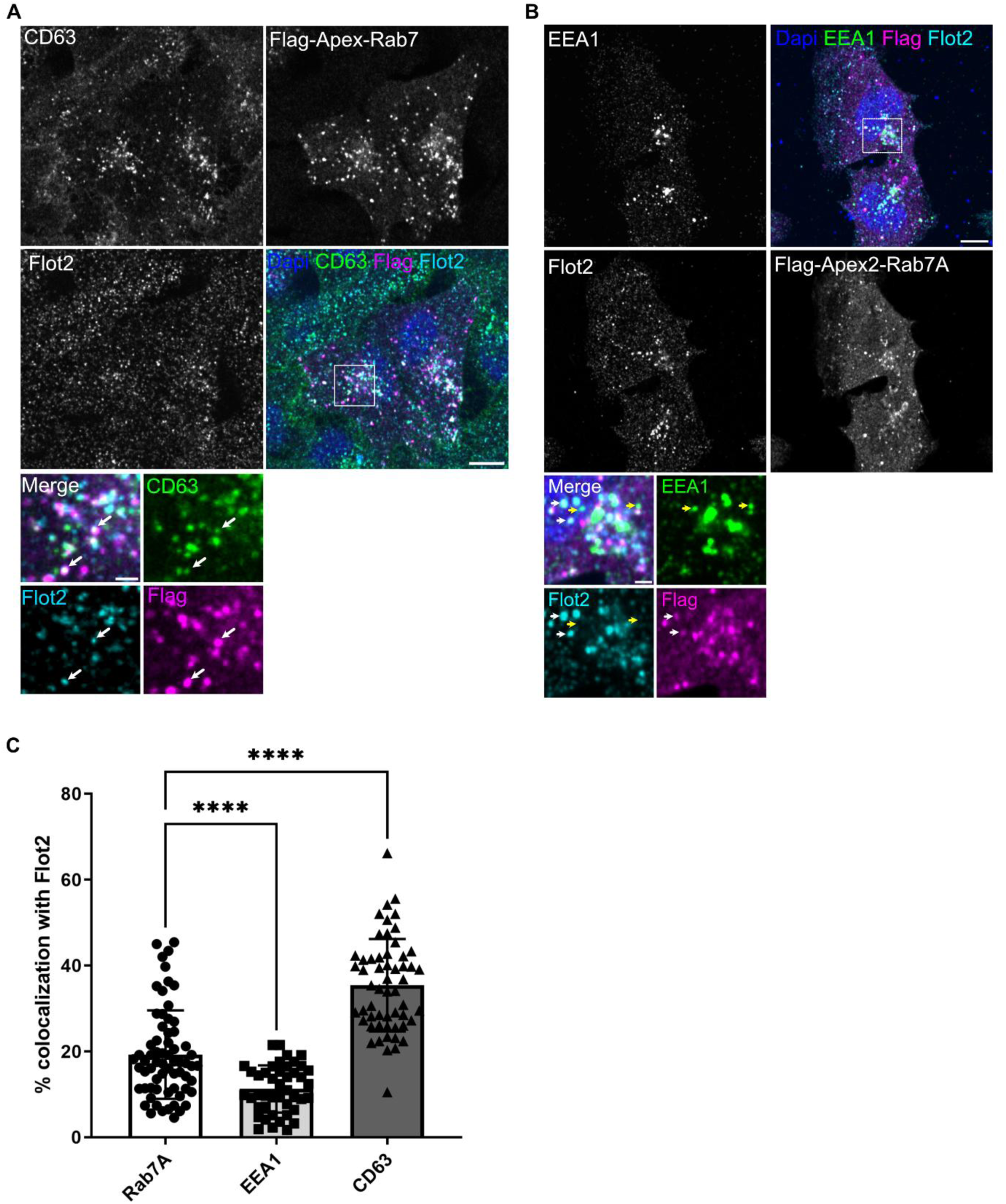
Flot 2 colocalizes with CD63 but less with Rab7A. (A) CD63 localizes with Flot2. SUM159 cells were transfected with Flag-Apex2-Rab7A. At 24 h post transfection, cells were visualized by confocal microscopy. Representative images from a single z-stack indicated CD63 (green), Flot2 (cyan) and Rab7A (Flag, magenta) using the anti-Flag antibody. White arrows indicate colocalization between Flot2, Rab7A and CD63. Scale bar = 10 µm and scale bar for zoomed regions=2µm. (B) EEA1 does not colocalizes with Flot2. SUM159 cells were transfected with Flag-Apex2-Rab7A. At 24 h post transfection, cells were visualized by confocal microscopy. Representative images from a single z-stack indicated EEA1 (green), Flot2 (cyan) and Rab7A (magenta) using the anti-Flag antibody. Scale bar = 10 µm and scale bar for zoomed regions = 2 µm. Colocalization with Flot2 and Rab7A (white arrows), no colocalization with EEA1 (yellow arrow). (C) The graph represents the percentage of colocalization of Flot2 with CD63, Flag-Apex2-Rab7A, and EEA1. Images from (A-B) were used. Data were plotted as percentage of colocalized volumes +/- s.e.m. Two-way anova test with multiple comparisons were used. ****p-value < 0.0001. Each dot represents 44 to 66 field of views from n = 3 independent experiments.

**Figure S4:**
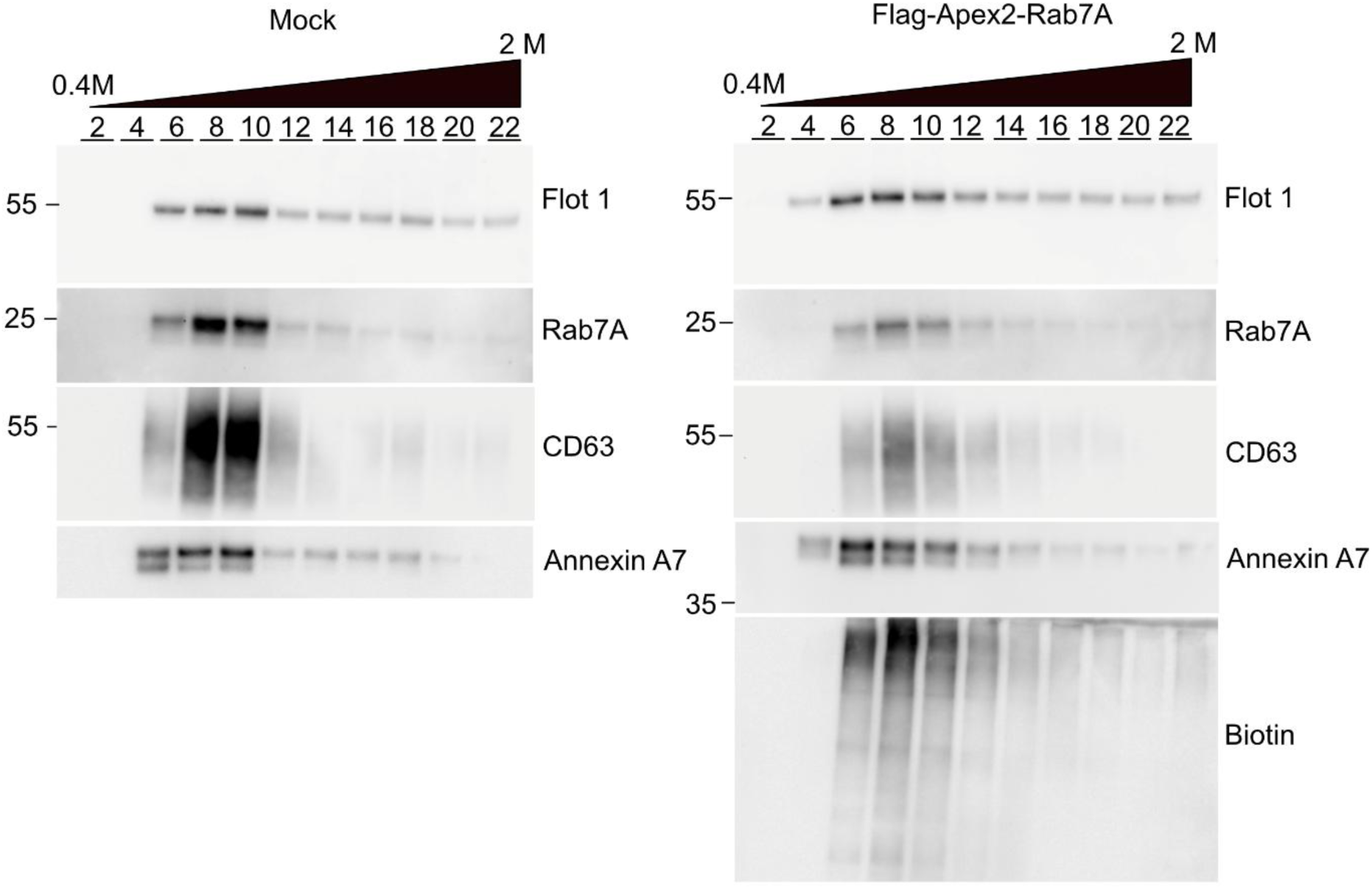
Density gradient performed on 100K EV pellet. EVs were collected and separated through differential ultracentrifugation. The 100K pellet is collected and layered on top of a continuous sucrose density gradient. Collected fractions were loaded from the lightest (0.4 M) to the heaviest density (2 M) on a gradient SDS-PAGE gel and western blot analysis were performed. Antibodies against Flot1, Rab7A, CD63, Annexin A7 and biotin were used.

**Figure S5:**
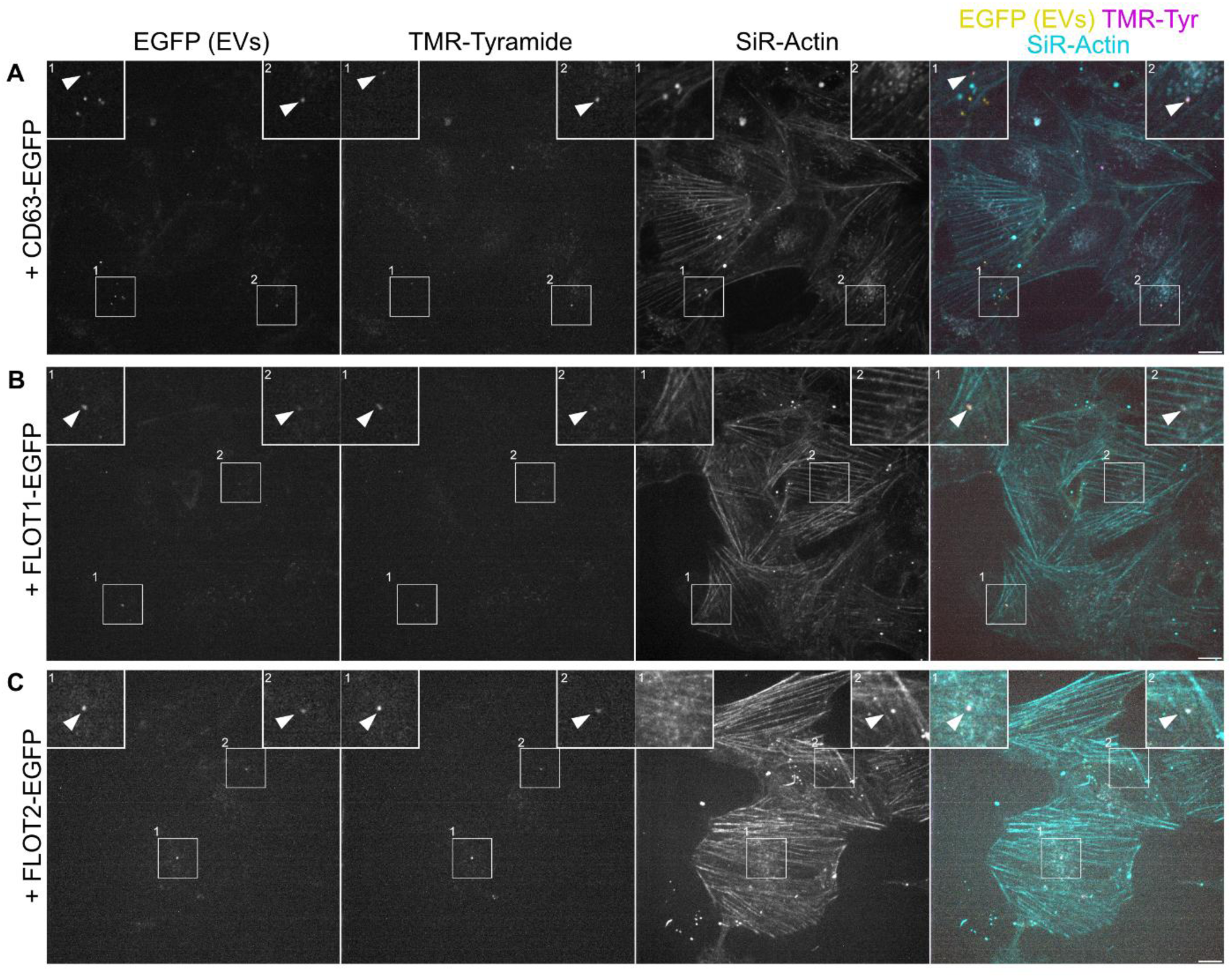
TMR-Tyramide labeled Flag-Apex2-Rab7A positive vesicles. (A-C) Representative confocal images from SUM159 wild-type cells treated with EVs from cell culture supernatants of donor cells co-expressing Flag-Apex2-Rab7A and CD63-EGFP (A), Flot 1-EGFP (B) or Flot 2-EGFP (C) (yellow) labelled with TMR-Tyramide (TMR-Tyr, magenta), isolated via differential ultracentrifugation. The surface of recipient cells was labelled using SiR-Actin (cyan). White arrowheads point to double-positive (EGFP+/TMR+) particles. Scale bar = 10 µm.

